# Pandemic-Scale Phylogenomics Reveals Elevated Recombination Rates in the SARS-CoV-2 Spike Region

**DOI:** 10.1101/2021.08.04.455157

**Authors:** Yatish Turkahia, Bryan Thornlow, Angie Hinrichs, Jakob McBroome, Nicolas Ayala, Cheng Ye, Nicola De Maio, David Haussler, Robert Lanfear, Russell Corbett-Detig

**Author notes:** Denotes equal contribution (Yatish Turkahia, Bryan Thornlow).

## Abstract

Accurate and timely detection of recombinant lineages is crucial for interpreting genetic variation, reconstructing epidemic spread, identifying selection and variants of interest, and accurately performing phylogenetic analyses. During the SARS-CoV-2 pandemic, genomic data generation has exceeded the capacities of existing analysis platforms, thereby crippling real-time analysis of viral recombination. Low SARS-CoV-2 mutation rates make detecting recombination difficult. Here, we develop and apply a novel phylogenomic method to exhaustively search a nearly comprehensive SARS-CoV-2 phylogeny for recombinant lineages. We investigate a 1.6M sample tree, and identify 606 recombination events. Approximately 2.7% of sequenced SARS-CoV-2 genomes have recombinant ancestry. Recombination breakpoints occur disproportionately in the Spike protein region. Our method empowers comprehensive real time tracking of viral recombination during the SARS-CoV-2 pandemic and beyond.

## Main Text

Recombination is a primary contributor of novel genetic variation in many prevalent pathogens, including *betacoronaviruses* (*1*), the clade that includes SARS-CoV-2. By mixing genetic material from diverse genomes, recombination can produce novel combinations of mutations that have potentially important phenotypic effects (*2*). For example, recombination is thought to have played an important role in the recent evolutionary histories of MERS (*3*) and SARS-CoV (*4*, *5*). Furthermore, a recombination event that transferred a portion of the Spike protein coding region into the ancestor of SARS-CoV-2 may have contributed to the emergence of the COVID-19 pandemic in human populations (*6*). Recombination is thought to have the potential to generate viruses with zoonotic potential in the future (*6*). Therefore, accurate and timely characterization of recombination is foundational for understanding the evolutionary biology and infectious potential of established and emerging pathogens in human, agricultural, and natural populations.

Now that substantial genetic diversity is present across SARS-CoV-2 populations (*7*) and co-infection with different SARS-CoV-2 variants has been known to sometimes occur (*8*), recombination is expected to be an important source of new genetic variation during the pandemic. Whether or not there is a detectable signal for recombination events in the SARS-CoV-2 genomes has been fiercely debated since the early days of the pandemic (*6*). Nonetheless, several apparently genuine recombinant lineages have been identified using *ad hoc* approaches (*9*) and semi-automated methods that cope with vast SARS-CoV-2 datasets by reducing the search space for possible pairs of recombinant ancestors (*9*, *10*). Because of the importance of timely and accurate surveillance of viral genetic variation during the ongoing SARS-CoV-2 pandemic, new approaches for detecting and characterizing recombinant haplotypes are needed to evaluate new variant genome sequences as quickly as they become available. Such rapid turnaround is essential for driving an informed and coordinated public health response to novel SARS-CoV-2 variants.

We developed a novel method for detecting recombination in pandemic-scale phylogenies, <u>R</u>ecombination <u>I</u>nference using <u>P</u>hylogenetic <u>PL</u>ac<u>E</u>ment<u>S</u> (RIPPLES, Fig. 1). Because recombination violates the central assumption of many phylogenetic methods, *i.e*., that a single evolutionary history is shared across the genome, recombinant lineages arising from diverse genomes will often be found on “long branches” which result from accommodating the divergent evolutionary histories of the two parental haplotypes (Fig. 1). RIPPLES exploits that signal by first identifying long branches on a comprehensive SARS-CoV-2 mutation-annotated tree (*11*, *12*). RIPPLES then exhaustively breaks the potential recombinant sequence into distinct segments that are differentiated by mutations on the recombinant sequence and separated by up to two breakpoints. For each set of breakpoints, RIPPLES places each of its corresponding segments using maximum parsimony to find the two parental nodes – hereafter termed donor and acceptor – that result in the highest parsimony score improvement relative to the original placement on the global phylogeny (Text S1). Our approach therefore leverages phylogenetic signals for each parental lineage as well as the spatial correlation or markers along the genome. We establish significance for the parsimony score improvement through a null model conditioned on the inferred site-specific rates of *de novo* mutation (Text S2-S3).

**Fig. 1.**
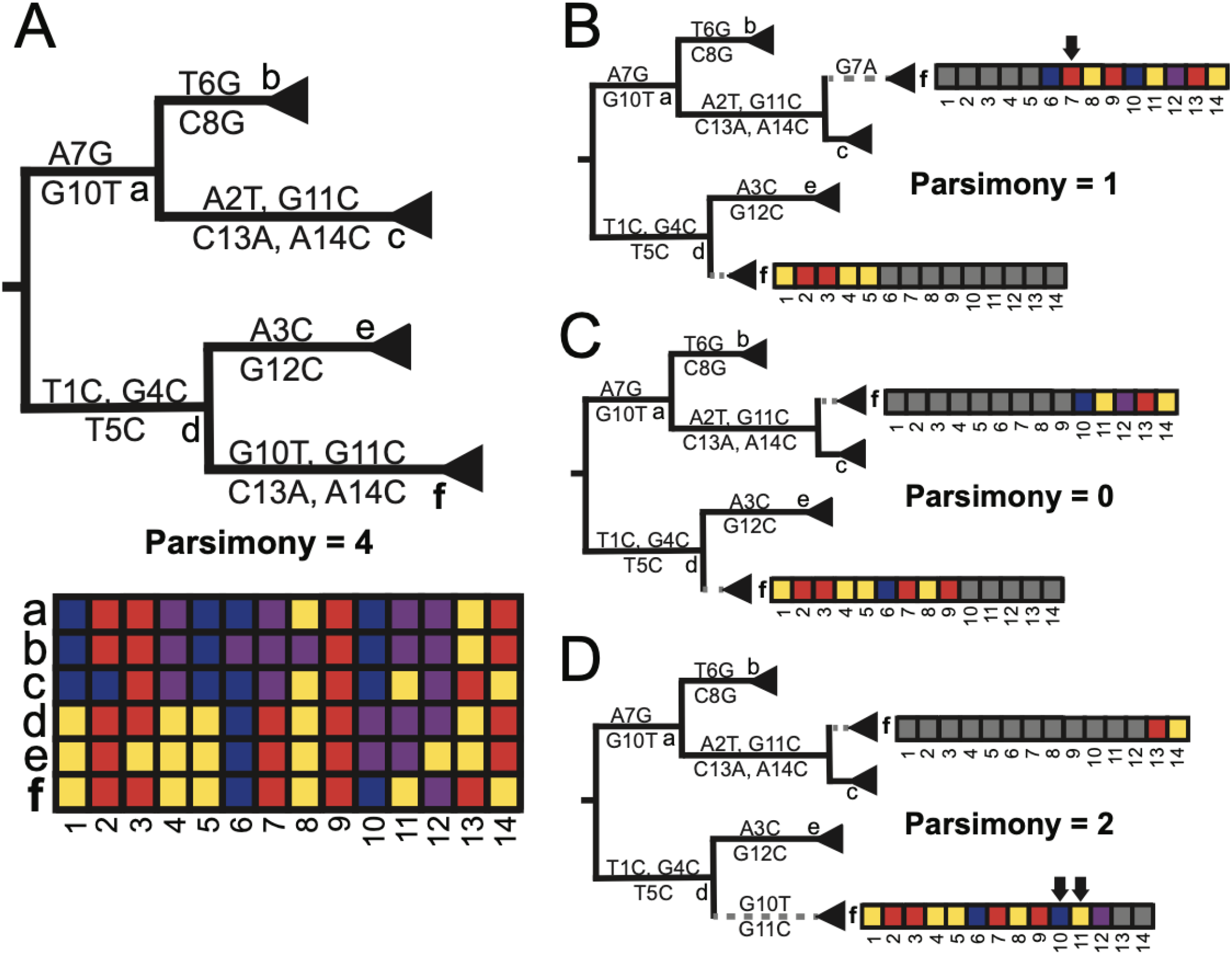
RIPPLES exhaustively searches for optimal parsimony improvements using partial interval placements. **(A)**: A phylogeny with 6 internal nodes (labeled a-f), in which node f is the one being currently investigated as a putative recombinant. The initial parsimony score of node f is 4, according to the multiple sequence alignment below the phylogeny, which displays the variation among samples and internal nodes. Note that internal nodes may not have corresponding sequences in reality, but test for recombination using reconstructed ancestral genomes. **(B-D)**: Three partial placements given breakpoints are shown with their resulting parsimony scores. Arrows mark sites that increase the sum parsimony of the two partial placements of f. The optimal partial placement and breakpoint prediction for node f is in the center (C), with one breakpoint after site 9 and with partial placements both as a sibling of node c and as a descendant of node d.

Substantial testing via simulation indicates that RIPPLES is sensitive and can confidently identify recombinant lineages. On our tree containing over 1.6 million SARS-CoV-2 sequences, RIPPLES takes just 6.25 minutes of wall time using 4 CPU threads and 1.94 GB of RAM per tested node, on average (Text S4-S5). Nonetheless, recombination breakpoints close to the edges of the SARS-CoV-2 genome are challenging to identify with certainty (Fig S1), which makes RIPPLES weakly biased against identifying recombination events near the edges of the viral genome. As expected, when recombination occurs between genetically similar sequences, it is harder to detect it using RIPPLES (Fig S2). The low identifiability of recombination events among closely related lineages is a well-known challenge in population genetics (*13*). Nonetheless, RIPPLES detects simulated recombinants with 93% sensitivity (Table S1), and is able to detect each of the highly confident recombinant samples identified by Jackson et al. (*9*) (Text S6). In contrast to previous methods for investigating recombination in the vast SARS-CoV-2 genomic datasets (*9*, *10*, *14*), RIPPLES can search for recombination on the inferred internal nodes of the phylogeny and does not require that phylogenetically informative sites or the set of parental lineages be selected *a priori*. This allows RIPPLES to achieve high sensitivity and be able to identify recombinant lineages without retraining the underlying model.

Recombination analysis using RIPPLES on a global phylogeny of approximately 1.6 million SARS-CoV-2 genomes reveals that a significant fraction of the sequenced SARS-CoV-2 genomes belong to detectable recombinant lineages. To mitigate the impacts of sequencing and assembly errors, we exclude all nodes with only a single descendant, we applied conservative filters to remove potentially spurious samples from the recombinant sets flagged by RIPPLES, and we manually confirmed mutations in a subset of putatively recombinant samples using raw sequence read data (Text S7-S8, Table S2, Fig S3). After this, we retained 606 unique recombination events, which have a combined total of 43,163 descendant samples (Table S3). This means that approximately 2.7% of total sampled SARS-CoV-2 genomes are inferred to belong to detectable recombinant lineages. *Post hoc* statistical analysis yields an empirical false discovery rate estimate of 11.6% for our statistical thresholds (Text S9, Table S4). Additionally, excess similarity of geographic location and date metadata among the descendants of donor and acceptor nodes strongly supports the notion that the ancestors of recombinant genomes co-circulated within human populations (Text S10, Fig S4) – a prerequisite for recombination. Because recombination events that occur between genetically similar viral lineages are challenging to detect (Fig S2), ours is expected to be a potentially large underestimate of the overall frequency of recombination. As a result, the RIPPLES estimate is likely conservative with respect to the global frequency of recombination in the SARS-CoV-2 population.

RIPPLES uncovered a strikingly non-uniform distribution of recombination breakpoint positions across the SARS-CoV-2 genome, consistent with previous analyses in *betacoronaviruses (15, 16)*. In particular, there is an excess of recombination breakpoints towards the 3’ end of the SARS-CoV-2 genome relative to expectations based on random breakpoint positions (p < 1×10^-7^; permutation test; Text S11). Importantly, no such bias is apparent when we simulate recombination breakpoints following a uniform distribution (Fig S1, Text S6). Change-point analysis identifies an increase in the frequency of recombination breakpoints immediately 5’ of the Spike protein region (20,787 bp; Text S12). The rate of recombination breakpoints is approximately three times higher towards the 3’ of the change-point than the 5’ interval (Fig 2) – which is similar to the relative recombination rates in the genomes of other human coronaviruses (*16*).

**Fig. 2.**
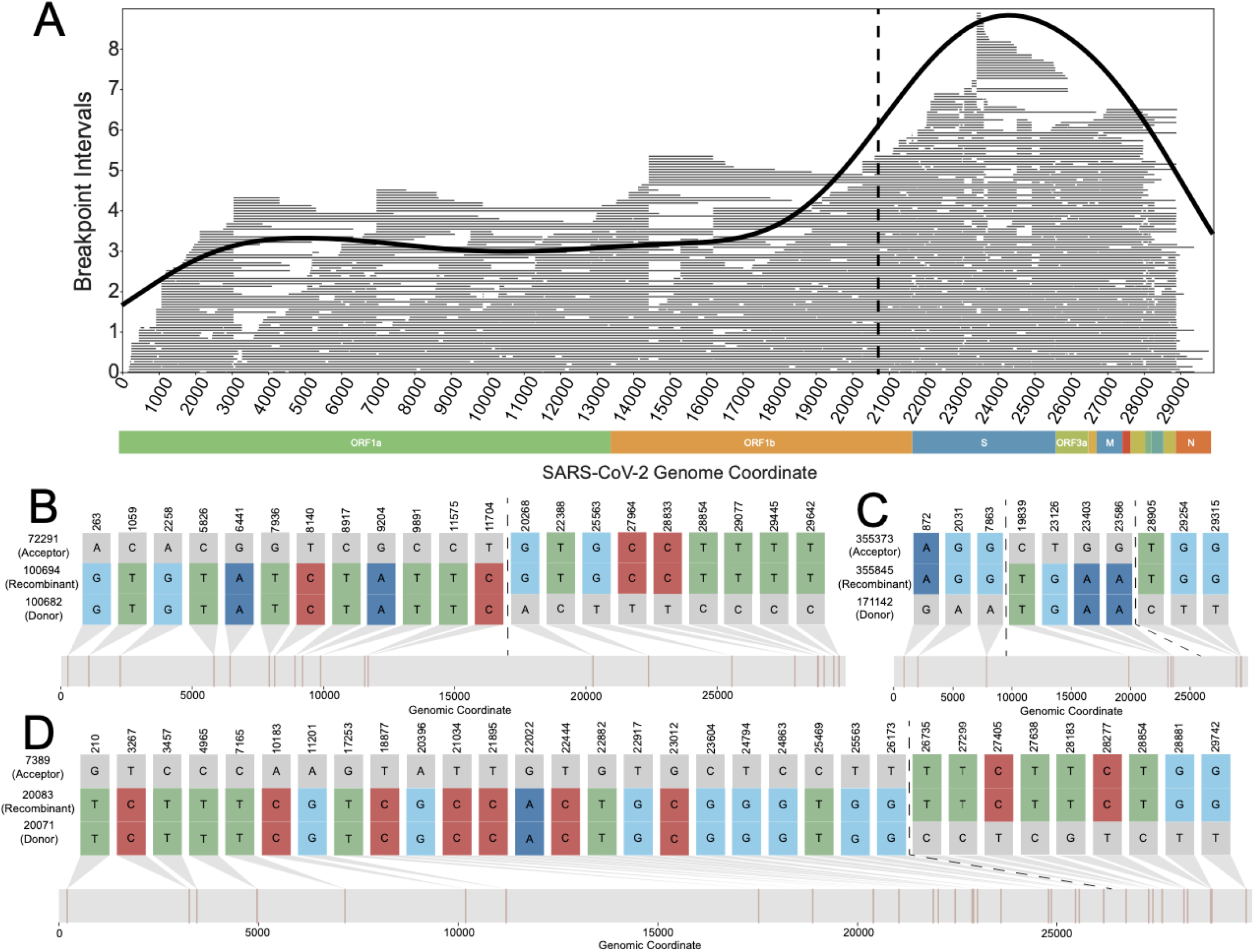
RIPPLES detects an excess of recombination in the Spike protein region. **(A)**: The distribution of midpoints of each breakpoint’s prediction interval are shown as a density plot, with the underlying recombination prediction intervals plotted as individual lines in gray. We used the midpoint of the breakpoint prediction interval because recombination events can only be localized to prediction intervals which are the regions between two recombination informative SNPs. A dashed vertical line at position 20,787 delimits recombination rate regions identified by change-point analysis (Text S12). The apparent lack of recombination towards the chromosome edges likely reflects a detection bias we describe above (Fig S2) **(B-D)**: Recombination-informative sites (i.e. positions where the recombinant node matches either but not both parent nodes) for three example recombinant trios detected by RIPPLES. The numbers to the left of each sequence correspond to the node identifiers from our MAT. B and D are examples of a recombinant with a single breakpoint (shown in dotted lines), C is an example of a recombinant with two breakpoints. B-D were generated using the SNIPIT package (https://github.com/aineniamh/snipit).

Several lines of evidence suggest that the skewed distribution of recombination breakpoint positions results primarily from a neutral mechanistic bias rather than being a consequence of positive selection. First, many of these recombinant clades have existed for a relatively short period of time, and might already be extinct. The mean timespan between the earliest and latest dates of observed descendants of detected recombinant nodes is just 37 days. Second, of the subset of recombination events that we inferred to occur between Variants of Concern (VOC; lineages B.1.1.7, B.1.351, B.1.617.2, and P.1 (*17*)) and other lineages, VOCs contribute slightly fewer Spike protein mutations than non-VOC lineages on average (58 out of 123 VOC/non-VOC recombinants, P = 0.765, sign test). Third, recombinant clade size does not greatly differ from the remaining clade sizes, which would be expected if recombinant lineages experienced strong selection (P = 0.8401, permutation test). Therefore, although natural selection on recombinant lineages could also impact the observed distribution of recombinant breakpoint positions (*16*), our data indicates that an important mechanistic bias shapes the distribution of recombination events across the SARS-CoV-2 genome.

Although not yet widespread among circulating SARS-CoV-2 genomes, recombination has measurably contributed to the genetic diversity within SARS-CoV-2 lineages. The ratio of variable positions contributed by recombination versus those resulting from *de novo* mutation, R/M, is commonly used to summarize the relative impacts of these two sources of variation (*15*). Using our dataset of putative recombination events, we estimate that R/M = 0.00264 in SARS-CoV-2 (Text S13). This is low for a coronavirus population (*e.g*. for MERS, R/M is estimated to be 0.25-0.31, (*15*)), which presumably reflects the extremely low genetic distances among possible recombinant ancestors during the earliest phases of the pandemic and the conservative nature of our approach. As SARS-CoV-2 populations accumulate genetic diversity and co-infect hosts with other species of viruses, recombination will play an increasingly large role in generating functional genetic diversity and this ratio could increase (*18*). RIPPLES is therefore poised to play a primary role in detecting novel recombinant lineages and quantifying their impacts on viral genomic diversity as the pandemic progresses.

Our extensively optimized implementation of RIPPLES allows it to search the entire phylogenetic tree and detect recombination both within and between SARS-CoV-2 lineages without *a priori* defining a set of lineages or clade-defining mutations. This is a key advantage of our approach relative to other methods that cope with the scale of SARS-CoV-2 datasets by reducing the search space for possible recombination events (*e.g., (9, 10, 14)*). RIPPLES discovers 239 recombination events within branches of the same Pango lineages. Our results also include 367 inter-lineage recombination events (Table S3). Additionally, we find evidence that recombination has influenced the Pangolin SARS-CoV-2 nomenclature system (*17*). Specifically, we discover that the root of B.1.355 lineage might have resulted from a recombination event between nodes belonging to the B.1.595 and B.1.371 lineages (Fig 3, Table S3). These diverse recombination events highlight the versatility and strengths of the approach taken in RIPPLES.

**Fig. 3.**
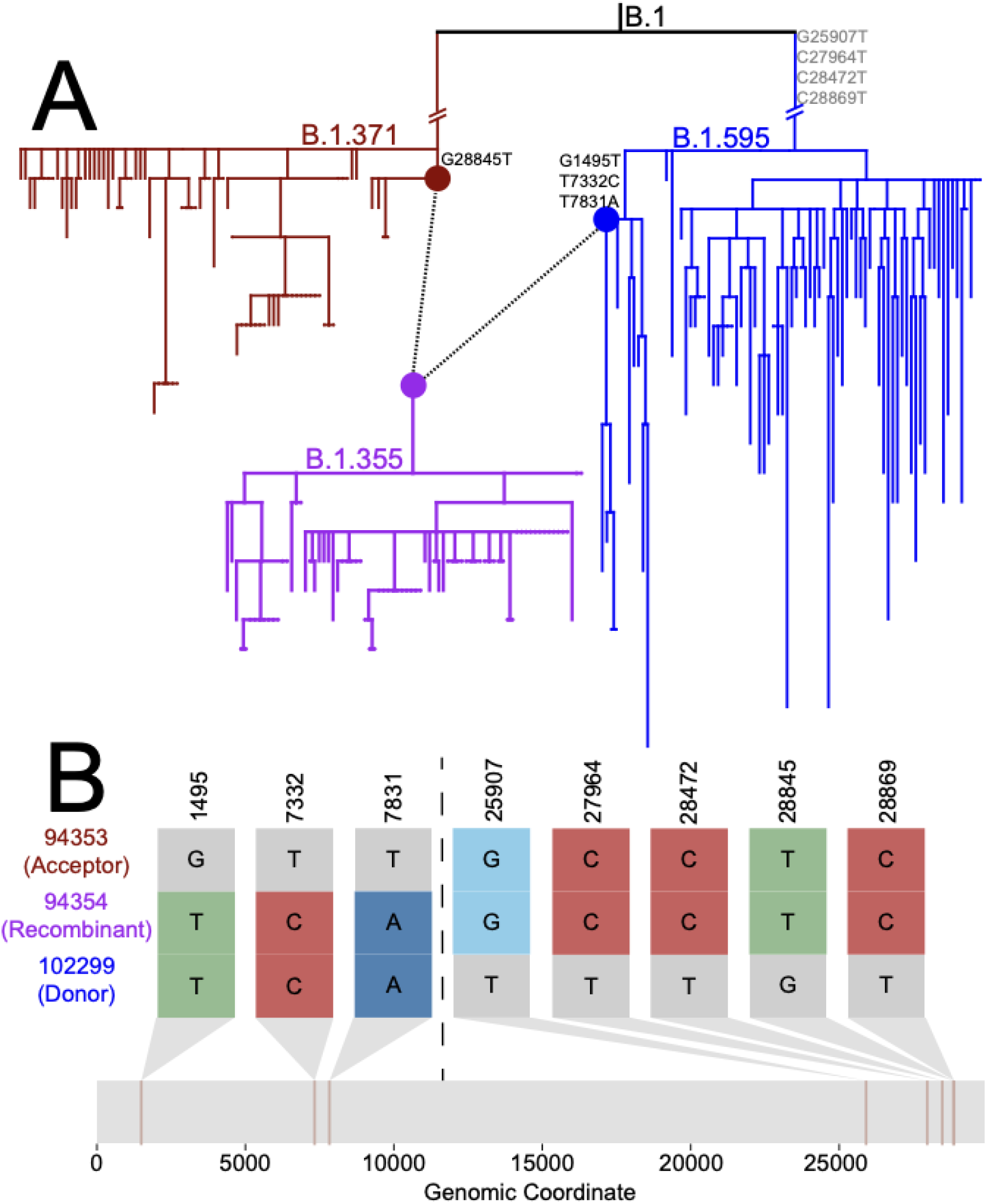
RIPPLES uncovered evidence that the B.1.355 lineage might have resulted from a recombination event between lineages of B.1.595 and B.1.371. **(A)**: Sub-phylogeny consisting of all 78 B.1.355 samples (purple) and the most closely related 78 samples to nodes 94353 and 102299 from lineages B.1.371 and B.1.595, respectively, using the “k nearest samples’’ function in matUtils (*12*). Nodes 94353 (red) and 102299 (blue) are connected by dotted lines to node 94354 (purple), the root of lineage B.1.355. Recombination-informative mutations are marked where they occur in the phylogeny, with those occurring in a parent but not shared by the recombinant sequence shown in gray. **(B)**: Recombination-informative sites (i.e. sites where the recombinant node matches either but not both parent nodes) are shown following the same format as Fig. 2B-D. B was generated using the SNIPIT package (https://github.com/aineniamh/snipit).

The detection of increased recombination rates around the SARS-CoV-2 Spike protein highlight the utility of ongoing surveillance. The Spike protein is a primary location of functional novelty for viral lineages as they adapt to transmission within and among human hosts. Our discovery of the excess of recombination events specifically around the Spike protein, as well as and the relatively high levels of recombinants currently in circulation, underline the importance of monitoring the evolution of new viral lineages that arise through mutation or recombination through real-time analyses of viral genomes. Our work also emphasizes the impact that explicitly considering phylogenetic networks will have for accurate interpretation of SARS-CoV-2 sequences (*16*).

Beyond SARS-CoV-2, recombination is a major evolutionary force driving viral and microbial adaptation. It can drive the spread of antibiotic resistance (*2*), drug resistance (*19*), and immunity and vaccine escape (*20*). Identification of recombination is an essential component of pathogen evolutionary analyses pipelines, since recombination can affect the quality of phylogenetic, transmission and phylodynamic inference (*21*). For these reasons, computational tools to detect microbial recombination have become very popular and important in recent years (*22*). The SARS-CoV-2 pandemic has driven an unprecedented surge of pathogen genome sequencing and data sharing, which has in turn highlighted some of the limitations of current software in investigating large genomic datasets (*23*). RIPPLES was built for pandemic-scale datasets and is sufficiently optimized to exhaustively search for recombination in one of the largest phylogenies ever inferred in just 89 hours on the n2d-highcpu-224 Google Cloud Platform (GCP) instance containing 224 vCPUs. To facilitate real-time analysis of recombination among tens of thousands of new SARS-CoV-2 sequences being generated by diverse research groups worldwide each day (*24*–*26*), RIPPLES provides an option to evaluate evidence for recombination ancestry in any user-supplied samples within minutes (Text S14). RIPPLES therefore opens the door for rapid analysis of recombination in heavily sampled and rapidly evolving pathogen populations, as well as providing a tool for real-time investigation of recombinants during a pandemic.

## Supporting information

Data S1: Phylogenetic tree used for this study, in newick format.

## Acknowledgments

We gratefully acknowledge the following Authors from the Originating laboratories responsible for obtaining the specimens, as well as the Submitting laboratories where the genome data were generated and shared via GISAID (Table S5) (*24*), China National Center for Bioinformation (Table S6), COVID-19 Genomics UK (COG-UK) (*26*) (Table S7), and National Center for Biotechnology Information database (*25*) (Table S8), on which this research is based. Additionally, the authors thank Shoh Mollenkamp for assisting with the code development.

## Funding

National Institutes of Health grant R35GM128932 (BT, JM, RC-D)

Alfred P. Sloan Foundation fellowship (RC-D)

National Institutes of Health grant T32HG008345 (BT, JM)

National Institutes of Health grant F31HG010584 (BT)

European Molecular Biology Laboratory (NDM)

Australian Research Council grant DP200103151 (RL)

Chan-Zuckerberg Initiative grant (RL)

University of California Office of the President Emergency COVID-19 Research Seed Funding Grant R00RG2456 (RC-D)

Additional funding provided by Eric and Wendy Schmidt by recommendation of the Schmidt Futures program (DH).

## Author contributions

Approach: RC-D, YT

Experimental design: RC-D, YT, BT, RL

Code development: YT, BT, AH, JM, NA, CY

Experiments: YT, BT, AH, NDM

Supervision: RC-D, DH

Writing: RC-D, YT

Editing: YT, BT, AH, JM, NA, CY, NDM, DH, RL, RC-D

## Competing interests

RL works as an advisor to GISAID. The remaining authors declare no competing interests.

## Data and Materials Availability

All data is available in the manuscript or the supplementary materials. RIPPLES software is available under the MIT license as part of the UShER package at https://github.com/yatisht/usher. We distribute RIPPLES with UShER because it uses the same underlying data objects and UShER is required to infer the input MAT. Documentation for RIPPLES and associated utilities can be found at https://usher-wiki.readthedocs.io/en/latest/.

## Supplementary Materials

### Supplementary Text

Text S1: Detection of Recombination with Partial Interval Placements

Text S2: Constructing a null model

Text S3: Establishing Significance Under A Null Model Based on Observed Mutation Rates

Text S4: Tree Optimization via Subtree Pruning and Regrafting (SPR) Moves

Text S5: Tree Pruning and Sample Filtration

Text S6: Establishing Sensitivity

Text S7: Filtering Possible False Positives

Text S8: Confirming Variation in Raw Sequence Read Datasets

Text S9: Empirical False Discovery Rate Estimation

Text S10: Measuring Spatial and Temporal Overlap of Recombinant Ancestors using Sample Metadata

Text S11: Permutation Test to Evaluate the Apparent Excess of 3’ Recombination

Text S12: Change-Point Analysis

Text S13: Estimating R/M

Text S14: Real time detection of recombinant ancestry in newly-sequenced samples

### Figures

Fig. S1: RIPPLES is highly sensitive and able to detect 93% of simulated breakpoints.

Fig. S2: RIPPLES more easily detects breakpoints causing large changes in parsimony score.

Fig. S3: Examples of detected trios filtered out due to sequence quality concerns.

Fig. S4: Recombinant Ancestors Exhibit Increased Spatial and Temporal Overlap.

### Tables

Table S1: Summary of simulated breakpoint detection.

Table S2: Raw sequence read datasets used to confirm recombination informative positions in selected recombinant samples.

Table S3: Summary of detected recombinant nodes.

Table S4: False discovery rate estimation for each parsimony score improvement observed in our dataset.

Table S5. Acknowledgements table for samples from GISAID.

Table S6. Acknowledgements table for samples from China National Center for Bioinformation.

Table S7. Acknowledgements table for samples from COVID-19 Genomics UK (COG-UK).

Table S8. Acknowledgements table for samples from INSDC Databases.

### Data

Data S1: Newick file containing phylogeny analyzed for recombination in this study.

**Fig. S1:**
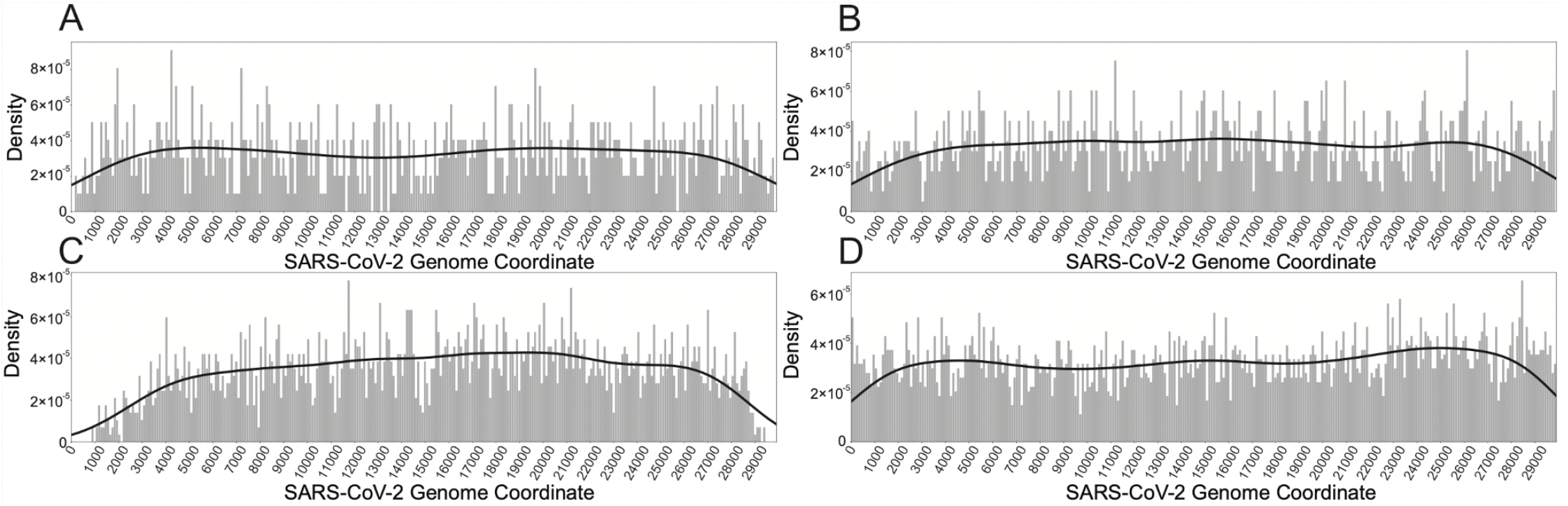
RIPPLES is highly sensitive and able to detect 93% of simulated breakpoints. We simulated 8,000 recombinant samples: 4,000 with one breakpoint and 4,000 with two breakpoints. The distributions of the true breakpoints for the simulated one- and two-breakpoint samples are shown in A and B, and were generated by choosing random positions across the genome, except that two breakpoints in a sample, if present, must be 1,000 nucleotides apart. The distribution of breakpoints detected for each simulated sample is shown by breakpoint position, with one-breakpoint samples in C, and two-breakpoint samples in D.

**Fig. S2:**
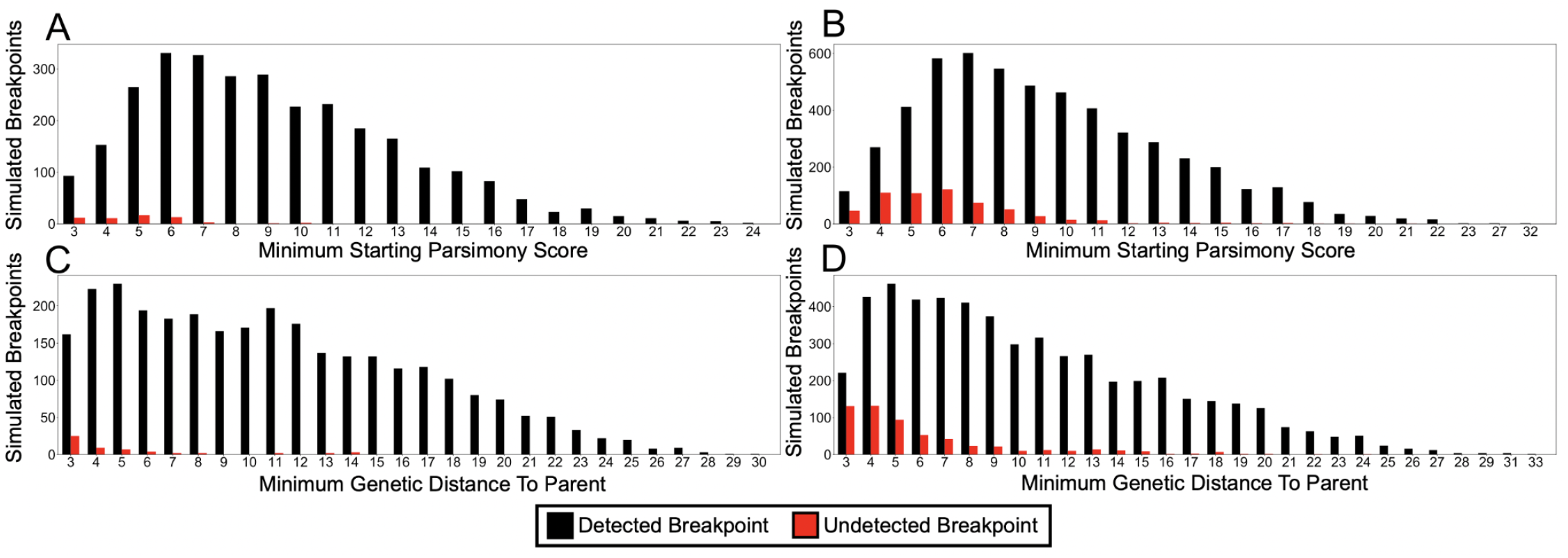
RIPPLES more easily detects breakpoints causing large changes in parsimony score. The distribution of simulated breakpoints detected for each simulated sample is shown by minimum starting parsimony score for the simulated one-breakpoint (A) and two-breakpoint (B) sample, and minimum genetic distance from simulated one-breakpoint (C) and two-breakpoint (D) sample to parent. Minimum starting parsimony (A, B) is dependent upon the initial placement of the recombinant node in the tree and refers to the genetic distance in mutations between the recombinant node and its direct parent in the phylogeny. Minimum genetic distance from sample to parent (C, D) refers to the number of mutations relevant to recombination that separate the recombinant node from either the donor or the acceptor, and is not dependent on the initial phylogeny. Detected breakpoints are shown in black and undetected breakpoints are shown in red. We condition on locating the true breakpoints and observing a significant parsimony score according to our phylogenetic null model. Therefore, we exclude recombination events with minimum starting parsimony scores and genetic distances of less than 3, as these are not significant under our null model.

**Fig. S3:**
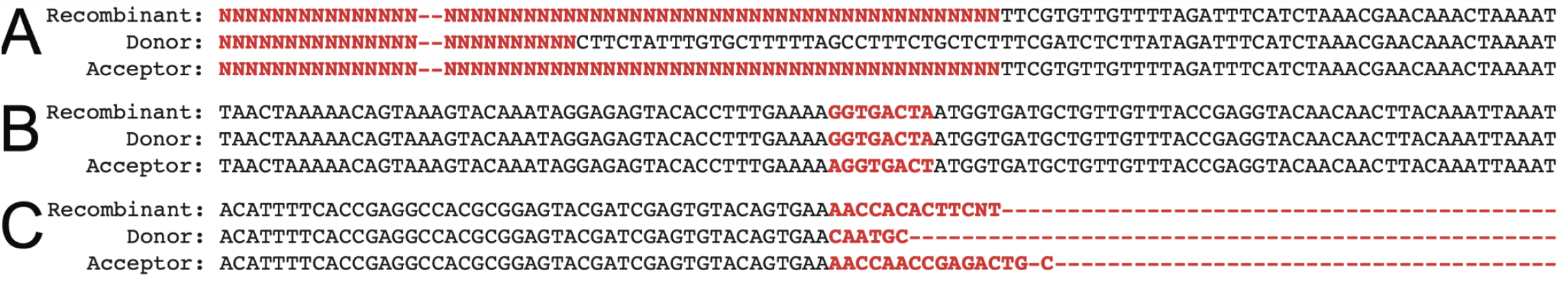
Examples of detected trios filtered out due to sequence quality concerns. **A:** Partial alignment of consensus sequences from a filtered recombinant trio of nodes 77695, 169585, and 77690, centered on site 28225, has consensus sequences of mostly ‘N’ spanning several sites meant to be informative of a recombination event. This can occur when many descendant samples have missing data. Mismatches between the three consensus sequences immediately flanking this region may be the result of poor sequencing quality as well. **B:** Partial alignment of consensus sequences from a filtered recombinant trio of nodes 173213, 173209, and 173274, centered on site 16846, has 7 recombination-informative mutations in an 8-nucleotide window that are unlikely to be true mutation events, but rather an alignment artifact or a complex indel event. **C:** Partial alignment of consensus sequences from a filtered recombinant trio of nodes 293461, 293460, and 211841, centered on site 29769, has 3 mismatches in a 5-nucleotide window, immediately flanked by a large gap in the alignment and are unlikely to be true mutations.

**Fig. S4:**
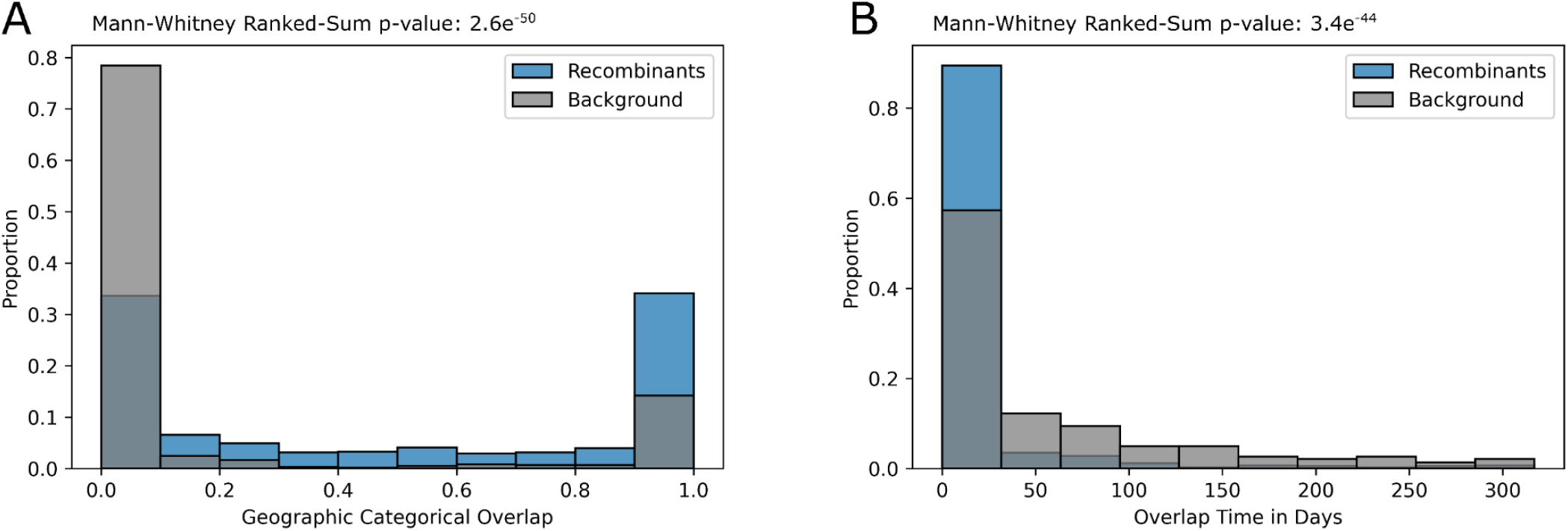
Recombinant Ancestors Exhibit Increased Spatial and Temporal Overlap. **A)** Spatial and **B)** temporal overlap for our recombinant trios (in blue) and the null distribution (in gray), with Mann-Whitney Ranked-Sum p-values for the statistical increase in overlap for the recombinant ancestors shown on the top.

## Supplemental Text

**Text S1. Detection of Recombination with Partial Interval Placements**

RIPPLES uses the space-efficient data structure of mntation-annotated trees (MATs) (*19*), in which the branches of the phylogenetic tree are annotated with mutations that have been inferred to have occurred on them, to identify recombination events. Figure 1 illustrates the underlying algorithm. RIPPLES starts by identifying putative recombinant nodes in the MAT which contain equal or more mutations on its corresponding branch than a user-specified value (the default value is 3). Next, for each putative recombinant node, RIPPLES infers the set of mutations that have occurred on its corresponding sequence by taking into account all mutations annotated on the branches on the path from the node to the phylogenetic root. To determine if this sequence is recombinant, RIPPLES then adds one or two breakpoints on the mutation sites to divide the sequence into two or three segments, respectively. The second segment is referred to as the donor segment, and the first and the third (in case of two breakpoints) segments are referred to as acceptor segments. RIPPLES then uses UShER’s (*18*) highly-optimized and multi-threaded phylogenetic placement module to evaluate the parsimony score of partially placing donor and acceptor segments, masking the mutations outside the segment boundaries, on every node of the phylogenetic tree, excluding the nodes that are direct descendants of the recombinant node.

The phylogenetic placement module permits breaking up an internal branch (placing part of the mutations of the branch above the breakpoint and the remaining below the breakpoint) to perform placement on a branch breakpoint if it results in a lower parsimony score. Next, RIPPLES stores all potential donor and acceptor nodes whose partial placement parsimony score is lower than the starting parsimony of the recombinant node. RIPPLES then limits the putative donor and acceptor lists to a maximum of 1000 nodes that provide the largest improvement in the parsimony score during the partial placement. This is to prevent the number of donor-acceptor pairs to become unmanageably large. RIPPLES evaluates each donor-acceptor pair to determine if the parsimony score of partially placing individual segments on the donor and acceptor nodes is lower than the parsimony score of the putative recombinant node by equal or above a user-specified threshold (the default is 3). If found, the putative recombinant node is flagged as a recombinant sequence of the donor and acceptor nodes. RIPPLES also takes into account the mutations on the donor and acceptor nodes to report the maximal genomic intervals within which breakpoint(s) could have occurred without increasing the parsimony score of the partial placements. RIPPLES repeats this process for all recombinant nodes and all possible breakpoint(s) within those nodes.

Because of some rare sub-optimality in tree structure, we sometimes notice that placing the whole putative recombinant node sequence solely on the donor or acceptor sequence can lower the parsimony score relative to the original placement. In such cases, RIPPLES measures the parsimony score improvement of the partial placements relative to the placement which provides the smallest parsimony score for the complete sequence.

**Text S2. Constructing a Null Model**

It is necessary to define a null model in order to determine whether we observe more recombination events than would be expected as false positives. Here, as an alternative to recombination, we define a null model wherein the additional mutations on a branch that we will test for recombination result instead from the underlying observed mutation process. To do this, we selected nodes at random and added k additional mutations, where k is an input parameter. Here, each mutation was drawn proportionally to the parsimony score of that mutation in the global phylogeny. This should make the extended branches we consider here consistent with the underlying null model. Importantly, our correction for *de novo* mutations should be more appropriate than alternative null models that assume that the mutation rate is equal across all sites (*e.g., (17)*). Furthermore, to whatever extent recombination contributes to apparently recurrent mutations, this model will be conservative for establishing significance under the null (below).

After generating sequences with additional mutations as described above, we placed those samples onto the phylogeny using UShER (*18*). Then, we searched for all possible partial placements using RIPPLES. We record the resulting improvement in parsimony score in the best partial placement that we found relative to the initial placement. The distribution of parsimony score improvement for each initial parsimony score provides a null model for the amount of improvement that might be expected under a model where mutation generates the long branches we search for and conditional on the phylogeny and the initial parsimony score.

**Text S3. Establishing Significance Under A Null Model Based on Observed Mutation Rates**

For each putative recombinant, we use the null distribution based on mutation on a single phylogeny without recombination to establish significance. For each node with a given initial parsimony score, we obtain the p-value as the proportion of simulated null distribution samples with the same initial parsimony score where the recombinant parsimony score improved by an equal number or more mutations than in the putative recombinant sample. Because the parsimony score improvement distribution is discrete and relatively small in value, the p-values obtained will typically be conservative. Furthermore, our test statistic is defined as the best possible parsimony score improvement for a given set of partial placements for a single node. The number of tests performed should therefore be linear with respect to the number of potential recombinant nodes evaluated. This property will typically be appealing when applying a false discovery rate correction because many tests will be highly correlated among possible parent nodes due to the nodes’ proximity within the phylogenetic tree. This can be a problem with methods that are not phylogenetic, *e.g*., those that examine all possible trios for donor-acceptor-recombinant relationships (e.g., (*45*)). With such methods, in some cases, if two nodes are distinguished by a SNP that is not contained within a recombinant segment, two or more ancestral nodes can yield identical results. More generally, closely related trios will yield highly correlated results which can impose important challenges for multiple testing corrections.

**Text S4. Tree Optimization via Subtree Pruning and Regrafting (SPR) Moves**

Optimizing that starting tree for RIPPLES is important for accurately estimating the parsimony score improvement in the partial placements of the recombinant sequence. We found that existing tree optimization tools, such as IQ-Tree (*46*), do not provide adequate speed and memory efficiencies to handle the massive SARS-CoV-2 phylogenies. Consequently, we developed our own fast and memory-efficient program, called matOptimize, to optimize the parsimony score of the massive SARS-CoV-2 phylogenies. Briefly, matOptimize starts with an input tree and a corresponding VCF file to annotate a set of bases, referred to as the Fitch set, that optimize the total parsimony score of the tree using the Fitch algorithm (*47*). All optimizations performed here were done using matOptimize program of commit 2981fcf from https://github.com/yatisht/usher.

Then, matOptimize begins the first optimization round by identifying a set of source nodes which have, or are ancestors to nodes having, recurring or reversal mutations. For each of these source nodes, matOptimize calculates whether a subtree pruning and regrafting (SPR) move for the node within a user-specified radius improves the parsimony score using the incremental update method of Gladstein et al. (*48*). matOptimize parallelizes this step across source nodes, assuming all moves are independent. Next, matOptimize identifies which of the profitable SPR moves could conflict (a pair of SPR moves are conflicting if they can affect each other’s parsimony score), and applies the non-conflicting profitable moves, prioritizing moves that provide larger parsimony score improvement among conflicting moves, and re-estimates the Fitch sets and parsimony scores for the affected nodes. matOptimize then starts a new optimization round, but restricts the source nodes to those that were found to improve the parsimony score in the previous round or were within the user-specified radius of the moves that did improve the parsimony score in the previous round. matOptimize keeps performing optimization rounds in this manner until it cannot find any moves that further improve the parsimony score for the entire round.

The matOptimize program is available under the UShER package (https://github.com/yatisht/usher) but further details of this method, as well as relative performance to other methods, will appear in a future publication.

**Text S5. Tree Pruning and Sample Filtration**

In order to test our method and detect as many SARS-CoV-2 recombination events as possible, we required a large phylogeny encompassing the genetic diversity of the virus. At UCSC, we have been maintaining a daily-updated SARS-CoV-2 phylogeny of all GISAID (*42*), GenBank (*43*) and COG-UK (*44*) sequences using the script https://github.com/ucscGenomeBrowser/kent/blob/master/src/hg/utils/otto/sarscov2phylo/updatePublic.sh and the method described in (*18*, *19*). We started with our phylogeny dated 28/05/2021 containing a total of 1,807,630 sequences with a parsimony score of 1,772,324. We then used the corresponding VCF file and masked all known problematic sites (*49*) and pruned out samples with fewer than 28,000 non-N nucleotides at positions where the SARS-CoV-2 reference genome had a non-N nucleotide. We also pruned out all samples with 2 or more ambiguous (non-[ACGTN-]) nucleotides, and then iteratively removed all samples on branches with length greater than 30 using the -*b 30* flag in matUtils. After this, we ran matOptimize twice using an SPR radius of 10 and 40 in subsequent rounds, and using the masked VCF as an input. Following this, we again iteratively pruned out all samples on branches with length greater than 30. The final tree contains 1,607,799 samples, 1,967,136 nodes, and has a total parsimony score of 1,522,210.

**Text S6. Establishing Sensitivity**

To test RIPPLES’ sensitivity, we simulated 8,000 recombinant samples by choosing 2 random internal nodes from our phylogeny with at least 10 descendants and choosing breakpoints at random across the genome. We generated 1,000 simulations each for one and two breakpoints with 0, 1, 2, and 3 additional mutations added to the sequence after the recombination event. We ensured that any two breakpoints were at least 1,000 nucleotides apart. The distribution of breakpoints selected for this experiment is approximately uniform, with slight bias against the ends of the chromosomes caused by this 1,000-nucleotide condition.

We then measured the ability of RIPPLES to detect breakpoints as a function of the position of the breakpoint and the minimum genetic distance from the recombinant node to either parent. Overall, we detect 93% of all breakpoints across all of our simulations (Table S1; Fig. S1–S2). Scripts used to generate simulated recombinants are available at https://github.com/bpt26/recombination/.

In addition to simulations, we evaluated the sensitivity of RIPPLES by asking if it could detect each of the high-confidence recombinant SARS-CoV-2 clusters of Jackson et al. (*16*). Briefly, this work used the unique and highly divergent B.1.1.7 haplotype to detect putative recombination events. To do this, we ran RIPPLES while relaxing the requirement that each detected recombinant have a minimum of two descendants. We did this because several of the clusters identified in that work have only a single extant descendant. We found that all putative recombination events identified in that work are also discovered by RIPPLES.

**Text S7. Filtering Possible False Positives**

We applied several *post hoc* filters to remove putative recombinant nodes that may be false positives resulting from several possible sources of error. For each internal node from each trio (putative recombinant, donor, and acceptor nodes) that comprised a recombinant event, we downloaded the consensus genome sequence for the nearest descendants of each node, from COG-UK, GenBank, GISAID, and the China National Center for Bioinformatics. We then aligned the sequences of all descendants for each trio using MAFFT (*50*), focusing specifically on recombination-informative sites, i.e. where the allele of the recombinant node matched one parent node but not the other. From each set of descendants, we created a consensus sequence for the recombinant, donor, and acceptor nodes. We then compared these consensus sequences to determine whether the informative sites for recombination were likely to be true mutations, or alignment artifacts not captured by our initial VCF file.

If an insertion or a deletion (indel) in the alignment or a set of missing bases (Ns) spanned at least one recombination-informative site in at least one of the consensus sequences, or if an informative site was within 5 nucleotides of an indel or set of missing bases at least 5 nucleotides long, or if more than 5 informative sites were within 2 nucleotides of an indel or set of Ns of any length, we discarded the trio. From careful inspection of individual trios, variation fitting these criteria might be influenced by sequencing quality (e.g. Fig. S3A). We also discarded trios containing more than 5 recombination-informative mutations in a 20-nucleotide span. While multi-nucleotide mutation events do occur, we found upon inspection of the raw sequences that cases of more than 5 mutations in such a small window most often occurred very near to either end of the sequence for that sample (Fig. S3B). We then discarded trios where 3 or more recombination-informative mutations in a 20-nucleotide span were found within 5 nucleotides of an indel or set of Ns at least 10 nucleotides in length (Fig. S3C). Finally, we removed trios for which the entire set of recombination-informative mutations in the donor or acceptor sequence occurred in a 20-nucleotide span. We have aimed to be conservative with our filtering and excluding these trios may eliminate some true variation from our dataset, but this conservative approach should limit false positives.

To further remove low-quality recombination events, we removed cases whose p-value in 3seq (*45*) was greater than 0.2. 3seq conducts non-parametric tests for clustering in sequences of binary values. We generated binary sequences using the informative sites for each trio (“A” if the recombinant matched only the donor, “B” if the recombinant matched only the acceptor). Our choice for a p-value of 0.2 is based on visual inspection of binary sequences. For example, a sequence of “AAAABBB” is assigned a p-value of 0.143, and “AAABBB” is assigned a p-value of 0.2. Our intention with this filter is to remove obviously erroneous recombination events, but a recombination event between nodes with few total informative sites could certainly result in such a sequence. However, the sequences “BAAABBABBBBBBBA” and “ABBBAABAAAAAAAB” result in p-values of 0.275. Clustering in these sequences do not resemble what we expect from simple recombination events and might be the result of contamination or mixed infections.

After controlling for sequence quality, we compared each parsimony improvement to the phylogenetically informed null model described above. We retained only trios whose p-value was less than 0.05, where the p-value represents the proportion of null samples, with parsimony score improvements of at least that observed for the sample of interest, given the same initial parsimony score. We then needed to remove redundant trios from this set of statistically significant predicted recombinants. Several recombinant nodes had predicted recombination events with different sets of parents, and/or different predicted breakpoint intervals, but because multiple recombination events are extremely unlikely to have occurred at one node, we retained only one recombination event for each node. To break ties, we favored recombination events for which we predicted only one breakpoint. Then, we favored trios with fewer informative sites. These represent cases where the donor and acceptor have more similar sequences, and we expect that strains with more similar sequences would be more likely to be in the same place at the same time, as is required for recombination to take place. After this, we resolved the remaining ties by favoring the trio with the smaller 3seq p-value, larger predicted breakpoint interval, and greater sum of descendants of the donor and acceptor nodes. Finally, we found a few cases where two predicted recombinant nodes were the acceptor or donor of each other, and retained only one event for these cases. To accomplish this, we applied the same set of sequential tiebreakers described above. After applying these filters, we retained 606 unique putative recombinant nodes, which are parents to 43,163 unique descendant samples. Scripts used for filtering results as described here are available at https://github.com/bpt26/recombination.

**Text S8. Confirming Variation in Raw Sequence Read Datasets**

To confirm the quality of samples informing the putative recombinant nodes, we manually examined the raw sequence reads for 10 of these samples where we could confidently link the raw sequence read data to a given consensus genome. These raw sequence reads were retrieved from NCBI’s SRA database, and were aligned to the Wuhan-Hu-1 reference genome using minimap2 (*51*). We then used samtools (*52*) to convert the output of minimap2 into manageable bam files, and to create bam index files. The bam files were examined at the informative sites using IGV (*53*), and they were found to have high consensus (>90%) in each case. This indicates the putative recombinants are unlikely to result from spurious signal due to sample contamination.

**Text S9. Empirical False Discovery Rate Estimation**

To estimate the false discovery rate associated with our specific approach and statistical threshold selected, we computed a *post hoc* empirical false discovery rate. To do this, we obtained the number of internal nodes that we tested and which were associated with a given parsimony score. Then, for each initial parsimony score and parsimony score improvement, we obtained the expected number of internal nodes that would display that parsimony score improvement under the null model, i.e. as a consequence of mutational processes and in the absence of recombination. We estimate the false discovery rate as the ratio of expected nodes for a given initial and final parsimony score to the number of detected recombinant nodes with the same initial and final parsimony score. As would be expected, more modest parsimony score improvements are associated with a higher estimated false discovery rate (Table S3).

**Text S10. Measuring Spatial and Temporal Overlap of Recombinant Ancestors using Sample Metadata** We performed *post hoc* analysis using sample metadata to determine if the ancestors of the recombinant nodes had higher spatial or temporal overlap than expected by chance, as follows. We treated geography as a categorical variable at the country level. We computed geographic overlap as the joint probability of choosing a sample from the same country from the descendants of the donor and the acceptor nodes. For temporal overlap, we recorded intervals from the earliest to the most recent sample descended from the donor and acceptor, respectively, and calculated the minimum number of days separating the two intervals (with 0 for overlapping intervals).

We generated a null distribution for both categories by selecting, for each detected trio, two random internal nodes from the tree with a number of descendants equal to the real donor and acceptor respectively. We then calculated geographic and temporal overlap in the same way for this random set. This is a conservative null because the most closely related nodes are often the most similar with respect to spatio-temporal metadata and therefore, the hardest to detect for recombination. Additionally, because of uneven sequencing efforts both temporally and geographically, as well as the sometimes ambiguous choice of donor and acceptor (e.g., because a mutation that distinguishes two nodes is not contained in a recombination tract), we would not expect perfect overlap in these metadata categories. It is important to note that we break ties between donor/acceptor pairs entirely without considering descendant metadata which would invalidate this *post hoc* analysis. Nonetheless, as we anticipate for true recombinants, both metadata categories are strongly enriched for similarity among donor and acceptor nodes.

**Text S11. Permutation Test to Evaluate the Apparent Excess of 3’ Recombination**

We next sought to determine if identified recombination breakpoints are shifted towards the 3’ end of the genome. To do this, we performed a permutation test comparing the difference of the mean of the distribution of detected breakpoints when recombination breakpoints are simulated uniformly at random with the mean of the breakpoint position distribution in the true set. Briefly, this is accomplished by randomizing the set of breakpoint positions between two vectors of equivalent lengths to the simulated and real sets. The reported p-value is the proportion of such permutations where the difference between the mean position of the true and simulated vectors was greater than or equal to the observed difference in the true data. Importantly, because both distributions reflect subsets of recombination events that can be detected conditional on the landscape of genetic diversity and phylogeny of SARS-CoV-2, this is an improved null comparison than assuming a distribution, *e.g*., a uniform distribution.

**Text S12. Change-Point Analysis**

To identify intervals where the frequency of recombination breakpoint intervals differs along the genome, we performed change-point analysis. We do this using the changepoint R package (*54*), and fit a poisson model to the count of recombination prediction interval midpoints. Because we have reduced ability to detect recombination events towards the edges of the SARS-CoV-2 genome, we removed the first and last thousand basepairs of the genome from this analysis. We then compute the mean rate of recombination breakpoints within the intervals on either side of the identified change-point to estimate the fold increase in recombination rate in the 3’ portion of the genome.

**Text S13. Estimating R/M**

A central focus of much of microbial evolutionary analysis is distinguishing the relative contributions of recombination and mutation to patterns of variation. To estimate this ratio for SARS-CoV-2, we conservatively assume that RIPPLES successfully detects all recombination events that are present on the phylogeny. Then, the decrease in parsimony score associated with each detected recombination event is an estimate of the total variation that results from recombination. The contribution to the total mutations present in the viral population is then the parsimony score decrease multiplied by the number of descendant lineages of that recombinant node. This is the total number of observed mutations whose genealogies contain a recombination event. For lineages descendant of multiple recombinant nodes, we multiplied by the recombinant node with greater parsimony score improvement. If we subtract this value from the total number of mutations observed across the entire datasets, we obtain an estimate of the number of mutations whose histories are attributed in whole to mutational processes. The ratio of these two numbers is an estimate of R/M averaged across all samples that are included in our tree.

**Text S14. Real time detection of recombinant ancestry in newly-sequenced samples**

Tens of thousands of new SARS-CoV-2 sequences are being added to online databases every day (*42*–*44*). In order to facilitate the real time discovery of novel recombination events as well as the presence of known recombinant ancestry in these sequences, RIPPLES includes an option to restrict the search for recombinant nodes to a set of user-specified samples and their ancestors. This allows RIPPLES to be used in conjunction with UShER (*18*) to incorporate newly-sequenced SARS-CoV-2 samples in the comprehensive global phylogeny and identify among them those with a recombinant ancestry in real time. For example, on a server using 40-core Intel Xeon E7-4870 processor, incorporating 10 new samples in a mutation-annotated tree (MAT) of 1.607 million SARS-CoV-2 sequences takes 26 seconds using UShER, and using the output MAT to identify from the newly-incorporated samples those having a recombinant ancestry takes an additional 8 minutes 34 seconds using RIPPLES, on average. We have added a tutorial for RIPPLES to our wiki, which is available at https://usher-wiki.readthedocs.io/en/latest/tutorials.html.

